# Bayesian modeling of the impact of antibiotic resistance on the efficiency of MRSA decolonization

**DOI:** 10.1101/2023.01.25.525621

**Authors:** Fanni Ojala, Mohamad R. Abdul Sater, Loren G. Miller, James A. McKinnell, Susan S. Huang, Yonatan H. Grad, Pekka Marttinen

## Abstract

Methicillin-resistant *Staphylococcus aureus* (MRSA) is a major cause of morbidity and mortality and one of the most serious infectious disease threats. Colonization by MRSA increases the risk of infection and transmission, underscoring the importance of decolonization efforts. However, success of these decolonization protocols varies, raising the possibility that some MRSA strains may be more persistent than others. Here, we studied how the persistence of MRSA colonization correlates with genomic presence of antibiotic resistance genes. Our analysis using a Bayesian mixed effects survival model found that high-level resistance to mupirocin was strongly associated with failure of the decolonization protocol. However, we did not see a similar effect with resistance to chlorhexidine or other antibiotics. Including strain-specific random effects improved the predictive performance, indicating that some strain characteristics other than resistance also contributed to persistence. Study subject-specific random effects did not improve the model. Our results highlight the need to consider the properties of the colonizing MRSA strain when deciding which treatments to include in the decolonization protocol.

**Author summary:** Methicillin-resistant *Staphylococcus aureus* (MRSA) is responsible for a high burden of morbidity and mortality. MRSA colonization incurs risk of MRSA infection and transmission, highlighting the need for highly effective decolonization protocols. However, decolonization protocols have had mixed success. The extent to which this mixed success is attributable to MRSA strain variations and their resistance to antibiotics, including those like mupirocin that are commonly used for decolonization, versus study subject factors, has been unclear. Here, we characterized the effect of antibiotic resistance genes on the efficiency of an MRSA decolonization protocol. We found that mupirocin resistance and strain-specific effects were associated with reduced effectiveness of an MRSA decolonization protocol, but that resistance to other antibiotics, including purported chlorhexidine resistance genes, and subject-specific effects had no discernible impact on protocol success. Our results highlight the need to consider the properties of the colonizing MRSA strain when deciding which treatments to include in the decolonization protocol.

## Introduction

*Staphylococcus aureus* colonizes approximately 30% of the population [1] and is a leading cause of healthcare and community associated infections [2]. Healthcare-associated infections with MRSA are associated with higher mortality rates as well as increased cost and hospitalization duration compared to patients infected with methicillin-susceptible S. *aureus*. MRSA carriers have a higher predisposition for infection with a 35% risk of MRSA infection within one year following colonization [3–6]. The anterior nares are the main reservoir of *S. aureus*, and the skin, particularly the axilla and groin, and pharynx are also often colonized. The risk of infection is correlated with the extent of colonization, as determined by the number of body sites found to be colonized [7]. MRSA infections are most often caused by the colonizing strain [8]. Infection prevention and control strategies include reducing spread by preventing colonization of new individuals as well as decolonization of MRSA carriers.

Decolonization reduces carriage rates and subsequent infection by 30% [9–12]. However, the effectiveness of decolonization protocols varies. The extent to which this variation is due to the protocol, the features of the MRSA strains colonizing the study subjects, characteristics of the colonized individuals, and the interaction among these factors has been unclear. Moreover, most studies have lacked appropriate controls and/or have had limited sample sizes [13]; additionally, most studies of decolonization protocols have had limited if any analysis of the colonizing MRSA strains [14].

The CLEAR (Changing Lives by Eradicating Antibiotic Resistance) Trial is a randomized controlled clinical trial of MRSA carriers comparing hygiene education to education plus decolonization after hospital discharge. The intervention arm underwent repeated decolonization involving a 5-day decolonization regimen applied every other week for six months. The decolonization regimen involved chlorhexidine antiseptic for daily showering and twice-daily mouthwash plus twice-daily mupirocin (a topical antibiotic) treatment of bilateral nares. The education (control) arm received hygiene education alone. Body site samples were collected five times: at enrollment, at one, three- and six-months post-enrollment during the intervention phase, and at nine months, which was three months after the end of intervention. Swabs were obtained from multiple body sites including nares, throat, skin (axilla and groin), as well as accessible wounds, if present. While the trial demonstrated the benefit of the decolonization protocol with chlorhexidine and mupirocin, persistent colonization was noted in a subset of both trial arms [15]. The factors that contributed to persistent colonization were not addressed in the primary manuscript.

The goal for the current investigation was to model the association between antibiotic resistance genes and the persistence of MRSA strains during a decolonization protocol. To study this, we used whole genome sequencing of 3901 isolates from 880 study subjects from CLEAR who had completed all follow-up visits. We used a Bayesian statistical framework (BaeMBac software) to define persistent strains [16] and formulated a Bayesian mixed effects survival model [17], where the survival outcome was the clearance of MRSA during a given study interval, and the lack of clearance represented persistent colonization. Resistances to different antibiotics as predicted by *Mykrobe predictor* [18] were the covariates in the fixed effects, and the study subject- and strain-specific random effects were included to quantify the impact of other subject and strain related factors on clearance. Our approach was fully Bayesian, which allowed characterization of uncertainty of all quantities of interest and incorporation of prior knowledge [19].

## Materials and methods

### Data

See [9] for the details of the study protocol. In brief, subjects were selected for the study based on an MRSA positive culture within the hospitalization prior to enrollment (0-month). Isolates were collected from the study subjects at 1-month, 3-month, 6-month and 9-month visits from the start of the study. Per protocol, the decolonization regimen was stopped six months after discharge, and therefore data from the 9-month interval were excluded from the subsequent analysis. A resistance profile based on the presence or absence of resistance genes was estimated for each isolate using *Mykrobe predictor* [18], except for chlorhexidine for which BLAST [20] was used to detect the presence of *qac* genes as a marker for resistance.The mupirocin resistance is detected by *Mykrobe* using *mupA* and *mupB* genes and represents high-level resistance. Other information included the observation time, sequence type, hospital, study subject ID and body site of the swab. Subjects in the decolonization arm that did not fully adhere to the treatment were excluded. Contaminated isolates and isolates marked as MSSA were removed.

The MRSA positive isolates from a single study subject were divided into strains using the software BaeMBac [16], where a ‘strain’ is defined as a population of genetically closely related isolates. The software uses a Bayesian model based on the single nucleotide polymorphism (SNP) distance and time between consecutive visits to estimate the probability that a pair isolates collected from a study subject represent the same strain. The SNP distance of 45, estimated by BaeMBac using 10 percent of the education arm data, was used as a threshold (see S1 Fig) to divide the isolates from the same subject into strains. The MRSA isolates were primarily from ST5 (N=1337) and ST8 (N=1968) [16]; isolates from the remaining STs (N=533) were excluded because of the small number of samples. Most subjects were colonized with only one strain over the course of the study, but some were colonized with multiple strains (Table 1).

**Table 1.** Distribution of the number of strains colonizing a study subject.

| #Strains | #Subjects |  |
| --- | --- | --- |
|  | Decolonization | Education |
| 1 | 160 | 287 |
| 2 | 24 | 78 |
| 3 | 1 | 27 |
| 4 | 1 | 4 |

### Preprocessing

Our goal was to study the clearance of MRSA using survival models. For this purpose, we defined observations (*y_i_*, **x**_*i*_, *t_i_*) in our survival data as follows. One observation *i* corresponded to one study interval (between consecutive follow-up visits) from one subject such that the subject was colonized by MRSA in the beginning of the interval. If the study subject cleared MRSA carriage during a given interval (e.g., from 1-month visit to 3-month visit), then *y_i_* = 1, otherwise *y_i_* = 0. The vector **x**_*i*_ specified the characteristics of the strain in the beginning of the interval, and it included the vector of indicators for resistance to different antibiotics. The time *t_i_* was simply the length of the interval in months. We assessed clearance at 1-month, 3-month and 6-month visits. We denoted the starting visit by *v*_0_ of the interval of interest (for example the 1-month visit) and *v*_1_ the end visit (for example the 3-month visit).

The covariates **x**_*i*_ included the presence of genetic markers for resistance of the colonizing strain at *v*_0_ to the following antibiotics: ciprofloxacin, clindamycin, erythromycin, gentamicin, mupirocin, rifampicin, tetracycline, trimethoprim and chlorhexidine. Penicillin and methicillin resistance were excluded, as all isolates were expected to be resistant. Vancomycin and fusidic acid were also excluded, because there was no resistance to these antibiotics. Statistics of the survival data are given in Tables 2 and 3.

**Table 2.** Summary of the survival data.

|  | Decolonization | Education |
| --- | --- | --- |
| No. intervals (i.e., observations) | 270 | 911 |
| No. cleared intervals ($y = 1$) | 173 | 401 |
| No. subjects for the intervals | 186 | 396 |
| No. strains in the intervals | 215 | 540 |

**Table 3.** Summary of the resistance profiles in the survival data.

| Resistant to | Decolonization | Education |
| --- | --- | --- |
| Ciprofloxacin | 227 (84.1%) | 797 (87.5%) |
| Clindamycin | 103 (38.1%) | 394 (43.2%) |
| Erythromycin | 223 (82.6%) | 816 (89.6%) |
| Gentamicin | 22 (8.1%) | 72 (7.9%) |
| Mupirocin | 30 (11.1%) | 74 (8.1%) |
| Rifampicin | 5 (1.9%) | 31 (3.4%) |
| Tetracycline | 8 (3.0%) | 33 (3.6%) |
| Trimethoprim | 8 (3.0%) | 29 (3.2%) |
| Chlorhexidine* | 33 (12.2%) | 114 (12.5%) |

Number of observations (i.e., intervals) in the strain-specific survival data corresponding to different antibiotic resistance types. (\*resistance to chlorhexidine is not well characterized, here we used qac genes as an indicator)

The data were formulated for survival analysis in two ways, as illustrated in Fig 1. First, in a strain-specific analysis, clearance was defined such that at *v*_1_ the subject was not colonized by the strain at any site (see Fig 1b for illustration). Furthermore, in the strain-specific analysis there may have been multiple colonizing isolates at *v*_0_ belonging to the same strain, and the covariate corresponding to a certain resistance was defined as present if at least one of these isolates was resistant; in practice, the isolates of the same strain were so closely related that their predicted full resistance profiles were identical in 86% of cases. Second, in a site-specific analysis, clearance of a strain was defined as the absence of the strain at *v*_1_ on a body site of interest (either no strain was observed at *v*_1_ or a strain different from the one at *v*_0_ was observed). If no swab was collected on *v*_1_, the observation was excluded from the survival analysis, except if the MRSA-positive *v*_0_ swab was taken from wound, when it was considered cleared by *v_1_* (wound healed), see Fig 1c. The preprocessing pipeline is summarized in S2 Appendix.

**Fig 1.**
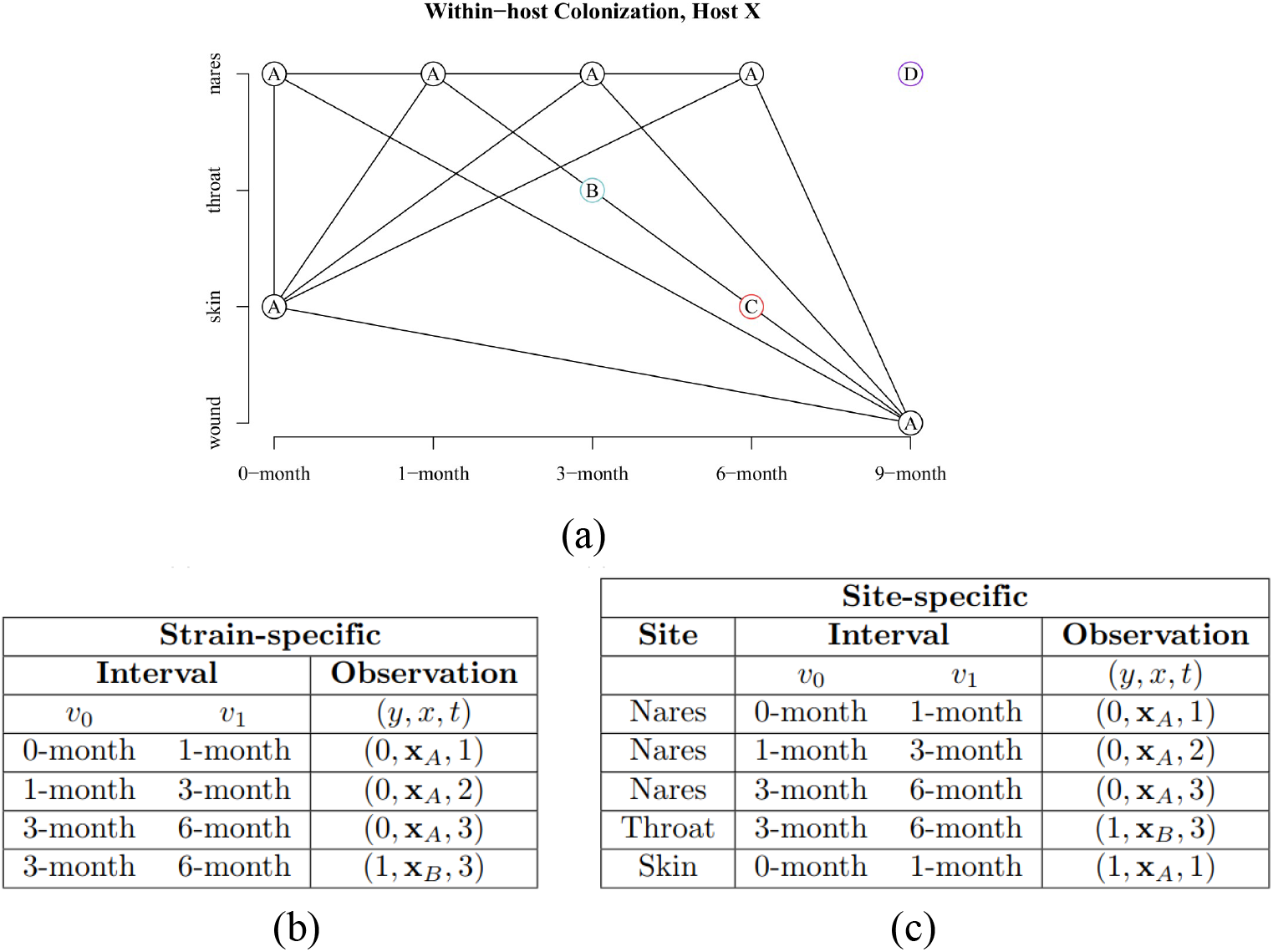
Strains in a multiply-colonized study subject. a) An example of a study subject colonized by four separate strains, A, B, C, and D, over the study period. b) Observations in the *strain-specific* survival data formulation for the subject. The subject contributed three intervals to the survival data, since the 9-month visit was excluded. Strains B and C were cleared immediately after acquisition, whereas strain A was persistent throughout the study. c) Observations in the *site-specific* survival data formulation (see text for details).

### Bayesian Survival Model

In survival analysis, the goal is to characterize time-to-event data in terms of the hazard of event or survival time until an event, affected by some covariates of interest. In our study, the fixed covariates were the presence or absence of resistance to each antibiotic, and the model included also subject- and strain-specific random effects. The parameters of interest were the fixed effect coefficients *β*, which denoted the magnitude of increase or decrease in risk or survival time for the covariates. Hence, formally, we were modeling the ‘hazard’ of clearance of an MRSA strain in a study interval from *v*_0_ to *v_1_* when the strain was known to be resistant to some antibiotics at *v*_0_. Consequently, an estimated hazard ratio *exp*(*β*) of 1.5, for example, indicated that a unit increase in the corresponding covariate resulted in a 1.5-fold risk of the clearance.

Our observations were either interval- or right censored: observations corresponding to study intervals where the clearance occurred (*y* = 1) between *v*_0_ and *v*_1_ were modeled as interval censored, as the exact event time of the clearance within the interval was not known. Right censoring was used for observations corresponding to study intervals in which the clearance event did not occur (*y* = 0) by the end of the interval *v*_1_.

The data consisted of

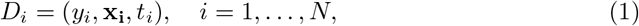

where *i* is an index for a visit interval such that *i* = 1, …,*N* comprised all visit intervals from all participants in the data set where the subject was colonized at the beginning of the interval, according to either the strain- or site-specific formulation, as described under the section *Preprocessing*. The response variable *y_i_* indicated whether the clearance happened within the interval, *t_i_* was the length of the interval, and vector x_i_ held the resistance profile at the beginning of the interval. The data in the decolonization and education arms were analysed separately. We used an exponential survival model with the proportional hazards parameterization. We assumed that the clearance rate was constant during a given interval. In addition, conditionally on the fixed covariates, the subject, the strain and the fact that the study subject was colonized in the beginning of the interval, the clearance probability in the interval was independent of clearances on other intervals (this assumption follows from the ‘memorylessness’ property of the exponential distribution). Hence, the hazard was given by

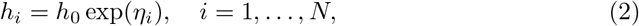

where *η_i_* was the linear predictor, defined as

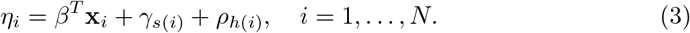

In Eq 3, *γ_s_*, *s* = 1, …, *S*, and *ρ_h_, h* = 1, …, *H*, were the the strain- and subject-specific random effects and *S* and *H* were the numbers of strains and study subjects, respectively. Functions *h*(*i*) and *s*(*i*) specified the subject (i.e. ‘host’) and the strain corresponding to interval *i*, respectively. The priors for the random effects were defined as

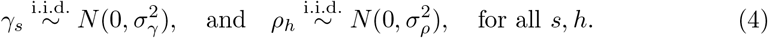

The survival function for interval *i* was thus

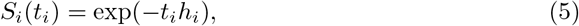

and it represented the probability that the study subject was still colonized by the same strain (i.e., “survived from clearance”) at the end of the interval. By letting *θ* denote jointly all model parameters, the log-likelihood function was defined as

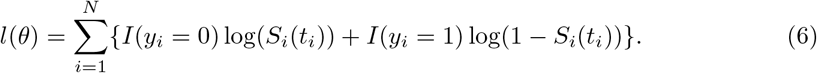

The priors for the coefficient *β* for the fixed effects and the intercept term *β*_0_ were defined to be relatively non-informative, i.e., to have a variance that exceeded the range of effects expected in the data, as follows:

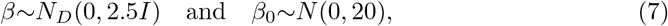

where *D* was the dimension of the fixed effects and *I* the identity matrix. The priors on the hyperparameters for the random effects (*σ_γ_* and *σ_ρ_*) were determined from the decomposition of the covariance matrix of the random effects into a correlation matrix Ω, a simplex *π*, and a scale parameter *τ*. Details of this decomposition can be found in *rstanarm* documentation and the Stan user guide [21, 22]. We set the hyperparameters as follows:

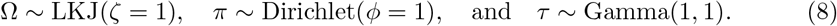

We estimated the posterior distributions for the parameters of interest by drawing samples from the posterior with an MCMC sampler, implemented using *rstanarm*’s function *stan_surv*. The R package *rstanarm* is an extension of the Stan programming language developed specifically as a platform for statistical analysis and Bayesian inference [21]. We ran the No-U-Turn sampler (a variant of Hamiltonian Monte Carlo) [23] for 7500 iterations for the strain-specific models and for 10000 iterations for the site-specific models over four chains. We increased the target average acceptance probability in the presence of divergent transitions as suggested by *rstanarm* documentation [24]. We assessed the convergence with 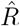 values and by visual inspection of traceplots. The full model included all antibiotics and both strain and study subject random effects. We compared the full model with different random effect configurations using the 10-fold cross-validation (CV). The coefficients *β* were estimated separately for the decolonization and education arms. The coefficient *β* represents the logarithm of the hazard ratio, which here means that the ‘instantaneous rate’ of clearance happening for a resistant strain is exp() times the rate for non-resistant strains.

## Results

### Exploratory data analysis

Before estimating the hazard ratios using the Bayesian approach, we conducted an exploratory analysis by calculating the clearance probability at *v*_1_ given resistance at *v*_0_ directly from the observation counts (intervals in the survival data) for resistant and non-resistant strains. This approach only considered one type of resistance at a time and neglected the different interval lengths. We saw that only 32% of the mupirocin-resistant observations were cleared at *v*_1_ in the decolonization arm while 67% of the non-resistant cases were cleared (Fig 2). The difference was significant with non-overlapping 95% confidence intervals. In the education arm, there was no difference in the clearance probability between resistant and non-resistant strains. Further, the clearance probability of the non-resistant strains is considerably larger in the decolonization arm than in the education arm, reflecting the overall efficiency of the protocol [9].

**Fig 2.**
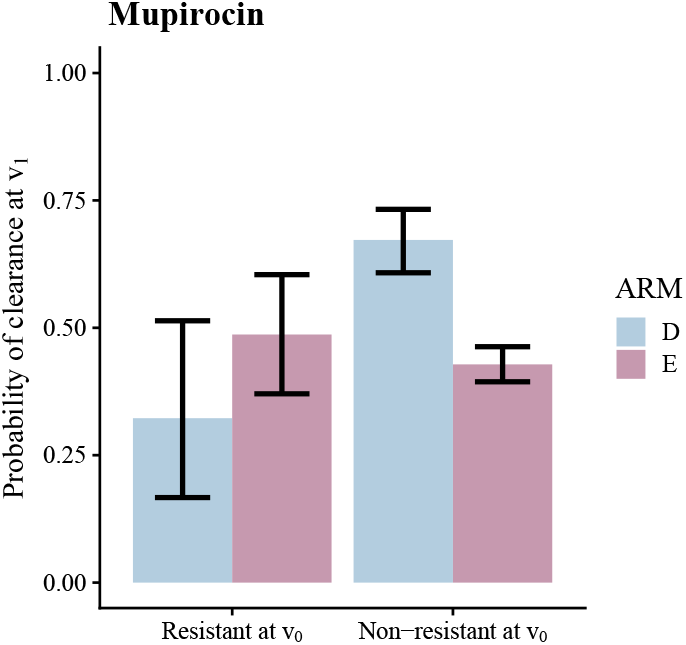
Clearance probabilities calculated from the counts of observations. Clearance probabilities given mupirocin resistance, computed directly from the counts of intervals colonized with resistant or non-resistant observations. On the y-axis, we have the clearance probability at the end of an interval, i.e., at *v*_1_, and the x-axis shows the resistance status at *v*_0_. The probability of clearance was calculated by dividing the numbers of persistent and cleared cases with the numbers of resistant or non-resistant observations in the data. The probability of clearance was lower for mupirocin-resistant strains than for non-resistant strains in the decolonization arm (D; blue). In the education arm (E; lavender), the probability of clearance (i.e., spontaneous loss of carriage) was the same regardless of the resistance status.

### Model comparison

We used the 10-fold cross-validation to compare the prediction accuracy of the different random effect combinations (no random effects, study subject, strain, study subject and strain). We quantified the results using the expected log-predictive density (elpd) [25], which is a metric for prediction accuracy. In both education and decolonization arms, including strain random effects improved the model considerably (Table 4). In contrast, including the subject-specific random effects did not improve the model, but instead slightly decreased the elpd value in the education arm. Because this decrease was minor and not significant, we decided to use the complete model to characterize both the strain and study subject random effects in the following section. The estimates for the fixed effects representing the impact of antibiotic resistance types on clearance were approximately the same regardless of whether the study subject random effects were included in the model or not (compare Fig 3 and S3 Fig).

**Table 4.** Model comparison.

| Model | Decolonization |  | Education |  |
| --- | --- | --- | --- | --- |
|  | elpd (mean) | elpd (SE) | elpd (mean) | elpd (SE) |
| Fixed | -208.0 | 9.6 | -689.1 | 12.9 |
| Fixed + Subject + Strain | -174.0 | 7.1 | -599.4 | 14.5 |
| Fixed + Subject | -174.1 | 7.5 | -671.6 | 14.9 |
| Fixed + Strain | -174.1 | 7.0 | -598.7 | 14.4 |

A model selection criterion, *expected log-predictive density*, measuring the prediction accuracy of the model, estimated with 10-fold cross-validation (mean and standard error) for different model formulations. Larger values of **elpd** indicate a better model.

**Fig 3.**
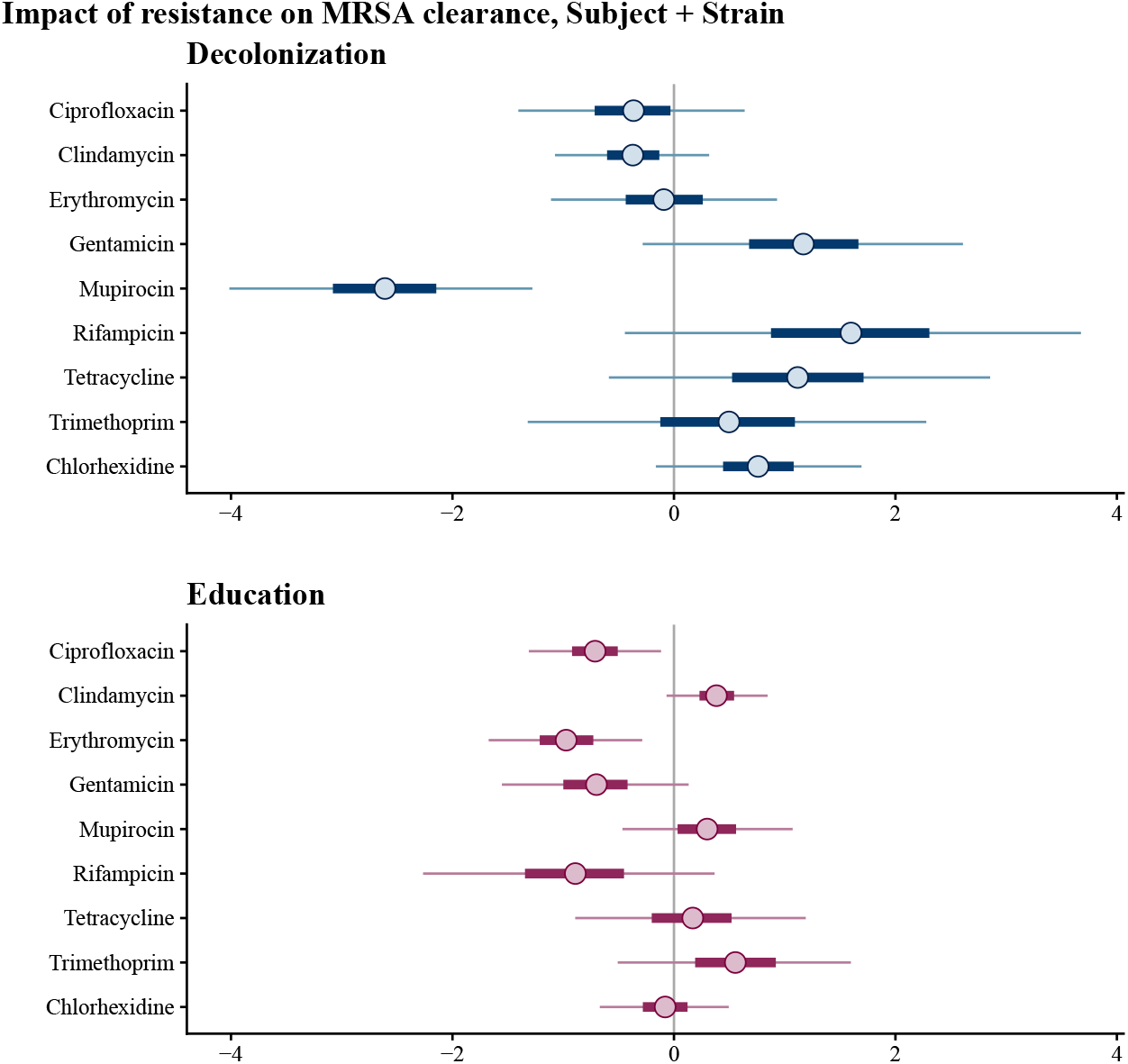
Credible intervals for the effects of antibiotic resistance types in the decolonization and education arms. 95% posterior credible intervals for the *β* parameters, representing the impact of each antibiotic resistance type on clearance. The model has study subject and strain random effects included and resistance types as fixed effects. A lower coefficient indicates a decreased rate (hazard) of clearance.

### Impact of antibiotic resistance on persistence

In the decolonization arm, the mupirocin resistance coefficient was -2.6 (95% CI is -4.0 to -1.3), indicating that the clearance rate of resistant strains was approximately 0.07 times the clearance rate of the non-resistant strains (Fig 3). Mupirocin resistance thus was correlated with greater MRSA persistence in the decolonization arm. However, this effect was not observed in the education arm (i.e., spontaneous loss was similar regardless of resistance). In contrast, chlorhexidine-resistant strains were not more persistent in the decolonization arm than the non-resistant strains, despite the use of chlorhexidine mouth and body-wash as part of the decolonization protocol. Furthermore, ciprofloxacin (–0.71, 95% CI: [-1.31, -0.12]) and erythromycin (–0.97, 95% CI: [-1.67, -0.29]) resistances were weakly associated with increased persistence in the education arm. Resistance to other antibiotics was not significantly associated with clearance, but the number of samples corresponding to some resistance types was limited (see Table 3), leading to wide credible intervals.

### Study subject and strain random effects

There was more variation in the strain random effects than in the study subject random effects in both the decolonization and education arms (Fig 4), which means that antibiotic resistance alone does not fully explain the variability in persistence. Furthermore, the variation in the strain random effects was larger in the education arm than in the decolonization arm. The study subject random effects were small in both arms. However, we note that many subjects were colonized by one strain only (see Table 1) and for those cases the effects of the strain and subject are statistically indistinguishable. Sequence type did not correlate with strain random effects (S4 Fig).

**Fig 4.**
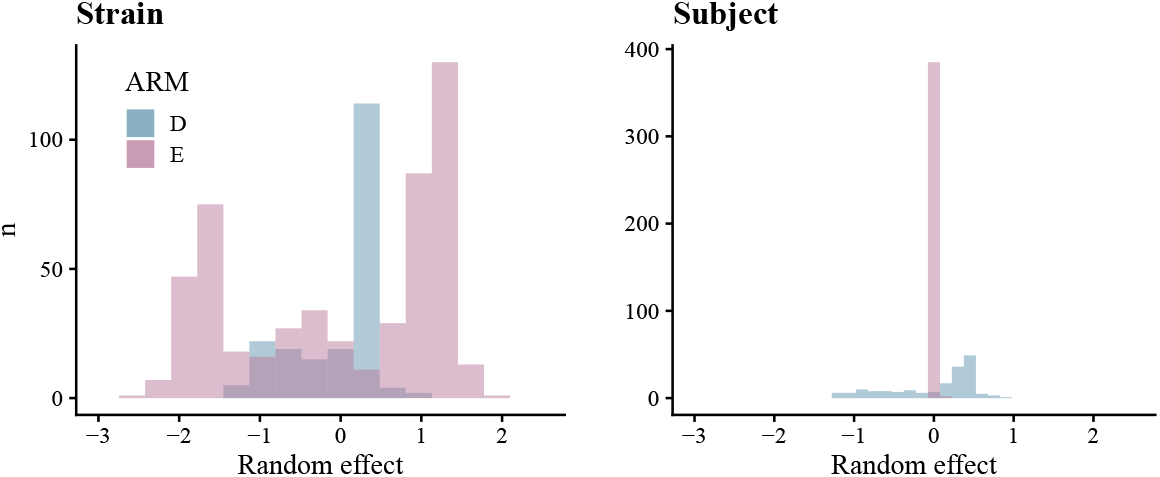
Estimated study subject and strain random effects. The figure shows histograms of the estimated strain and study subject-specific random effects. In both the decolonization (D) and education (E) arms, there was more variability in the strain random effects than in the study subject random effects.

### Site-specific analysis

In the decolonization arm, mupirocin resistance was again strongly associated with a reduced rate of clearance (i.e., increased persistence) in the nares (–2.26, 95% CI: [-3.8, -0.87]) but did not significantly correlate with clearance at other body sites (Fig 5, S5 Fig). Ciprofloxacin resistance and gentamicin resistance were weakly associated with increased persistence (–0.99, 95% CI: [-1.67, -0.32] and –1.12, [-2.06, -0.21]) in the nares in the education arm. In addition, we saw possible weak associations between chlorhexidine resistance and decreased persistence in the throat in the decolonization arm (2.07, 95% CI: [0.13, 4.00]), and between tetracycline resistance and increased persistence in the wound in the education arm (–1.65, 95% CI: [-3.32, -0.11]) (S5 Fig). The variation in the strain random effects was again greater than in the subject random effects (S6 Fig). Furthermore, this effect was clearest in the nares, from which most of the samples were obtained.

**Fig 5.**
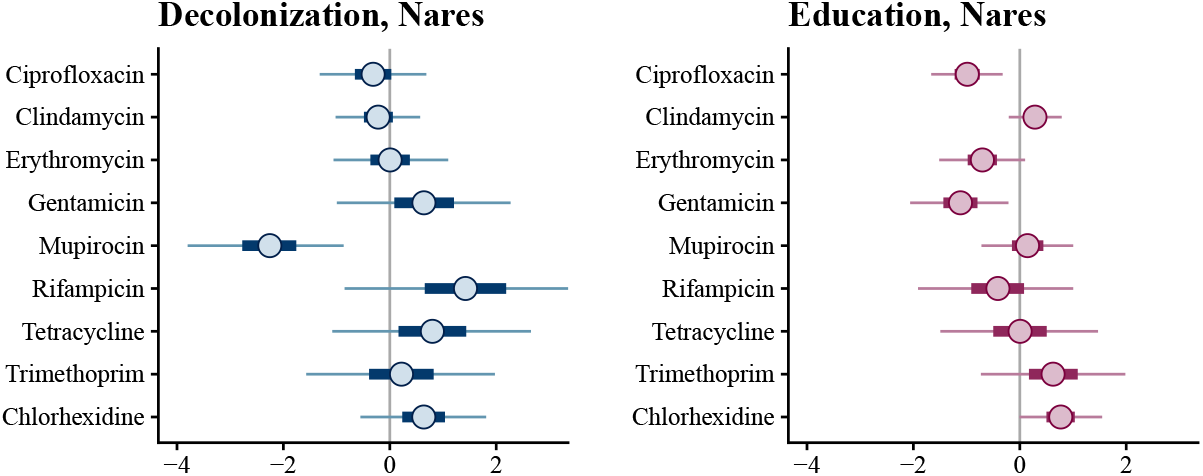
Results of the site-specific analysis. The figure shows 95% credible intervals for the effect of each antibiotic resistance type on clearance in both study arms. The results for the nares are shown here, as it had the largest effect, and for the other sites in S5 Fig

### Discussion

We applied Bayesian survival analysis on a dataset of sequenced MRSA samples collected from colonized patients after hospital discharge at given intervals during a follow-up period. Our results showed that mupirocin-resistant MRSA strains were more persistent than non-resistant strains in the decolonization arm, but not in the education only arm. When we looked at each body site separately, the effect of mupirocin was detected only in the nares, and not in the skin, throat, or wound. Since mupirocin is administered intranasally as part of the decolonization protocol and nares is the most prominent site of MRSA colonization [26], this result seems expected. However, despite chlorhexidine also being part of the decolonization protocol, chlorhexidine resistance did not seem to be associated with decolonization failure. This could be because chlorhexidine is applied to the throat (mouth wash), and skin and wound (baths), but not intranasally, but is most likely because genetic correlates of chlorhexidine resistance are quite poor. In the site-specific analysis, we saw a possible association between chlorhexidine and *decreased* persistence in the throat; however, the credible interval only slightly excluded zero, and hence this could be due to noise or otherwise reflect an unknown interaction between resistance types and persistence. Our study also confirmed the previously reported overall increased probability of clearance in the decolonization arm [9].

In the education arm, resistance to ciprofloxacin and erythromycin were found associated with increased persistence. A plausible biological mechanism is that some antibiotics had been used by the study subjects, giving resistant strains an advantage over non-resistant strains. A similar explanation could apply to the findings from the site-specific analysis, where we observed an increase in persistence in the education arm for gentamicin and ciprofloxacin-resistant strains in the nares and for tetracycline-resistant strains in wounds. Another possible explanation is that the order of causation is reversed: a strain that is persistent might be more likely to become antibiotic-resistant than a non-persistent strain, perhaps due to increased antibiotic exposure or shared biological mechanism. Further studies are needed to distinguish among these hypotheses.

Strain-specific random effects improved the ability of the model to predict persistence, which suggests that another genomic factor beyond the resistance determinants may affect persistence. The sequence type (ST5 vs ST8) of the MRSA strain did not seem to be associated with persistence S4 Fig. Including the study subject-specific random effects in addition to the strain-specific effects did not improve the model, in contrast with reports that study subject related factors affect MRSA colonization (for example the nasal microbiota [27]). However, we note that some strain and subject-specific effects were overlapping (when there was only one strain from one subject), and consequently some subject effects might be explained as part of the strain effects.

In future studies, a larger dataset could confirm associations between resistance and persistence that did not reach statistical significance in this study. This could also help to identify in more detail the genetic determinants of persistence that were represented here by the strain-specific random effects. While the random effects for study subjects were not significant in our study, including explicit characteristics of the subjects might add power to find some features that affect persistence. In addition, the use of antibiotics other than those that were part of the protocol during the study period was not considered in this study, but including them as covariates might reveal further insights about the relationship between resistance and persistence.

### Conclusion

We showed that genetic variants for mupirocin-resistance in MRSA were associated with a large drop in the efficiency of a decolonization protocol that includes intranasal mupirocin. Therefore, alternative decolonization protocols for patients with mupirocin-resistant MRSA colonization should be considered, such as nasal iodophor in place of mupirocin, although mupirocin is a superior treatment of the two in general [28]. However, we did not see a similar effect for chlorhexidine body and mouth wash, another part of the decolonization protocol, nor for any other antibiotic, which supports chlorhexidine as a reliable component of a skin decontamination protocol even when genetic correlates of chlorhexidine resistance are identified. In general, these findings point to the potential utility of improving the efficiency of decolonization protocols by characterizing an individual’s colonizing strain to determine its resistance profile.

## Supporting information

**S1 Fig. BaeMBac output for same strain probability** Contour plot detailing the same strain probability with SNP distance *d*\* on the x-axis and the time between consecutive visits (in generations) on the y-axis. The probability of 0.5 was used to decide the threshold distance of 45 SNPs that was used to classify a pair of MRSA isolates observed in consecutive visits as the same or different strain. The BaeMBas was run using 10 percent of randomly selected isolates from the education arm. The threshold was not sensitive to the amount of data, and the decolonization arm was not used as the BaeMBac software assumes a model of neutral evolution when calculating the same strain probability.

**S2 Appendix. Preprocessing pipeline** Preprocessing steps to format the data for the survival analysis. *n_g_* represents the number of genomes (isolates), *n_i_* the number of intervals and *n_s_* the number of study subjects with sequencing data for the colonizing strains available. The number of observations is smaller after “Restructure data”, because the survival data are considered by interval: recruitment to 1-month, 1-month to 3-month, 3-month to 6-month and 6-month to 9-month. For example, in the original data one body site could have an isolate at each of the visits, contributing in total five observations in the original data. However, we only have four intervals to consider in the case of survival data, because we are interested in the clearance status of an MRSA strain between or at the end interval of two consecutive isolates.

**S3 Fig. Estimated fixed effects with only strain random effects in the model** The 95% credible intervals for parameters *β* for all antibiotics with strain random effects included in the model, but excluding the subject-specific random effects.

**S4 Fig. Estimated strain random effects for sequence types ST5 and ST8.** The strain-specific survival model was used to estimate the strain random effects. Histograms show the distributions of the strain random effect posterior means in the decolonization and education arms.

**S5 Fig. Throat, skin and wound posterior intervals** 95% posterior intervals.

**S6 Fig. Estimated study subject and strain random effects by site.** Strain random effects have more variation in the posterior means than study subject random effects, most notably in the nares.

