## Supplementary figures and images for "Bayesian modeling of the impact of antibiotic resistance on the efficiency of MRSA decolonization"

### S1 Fig.

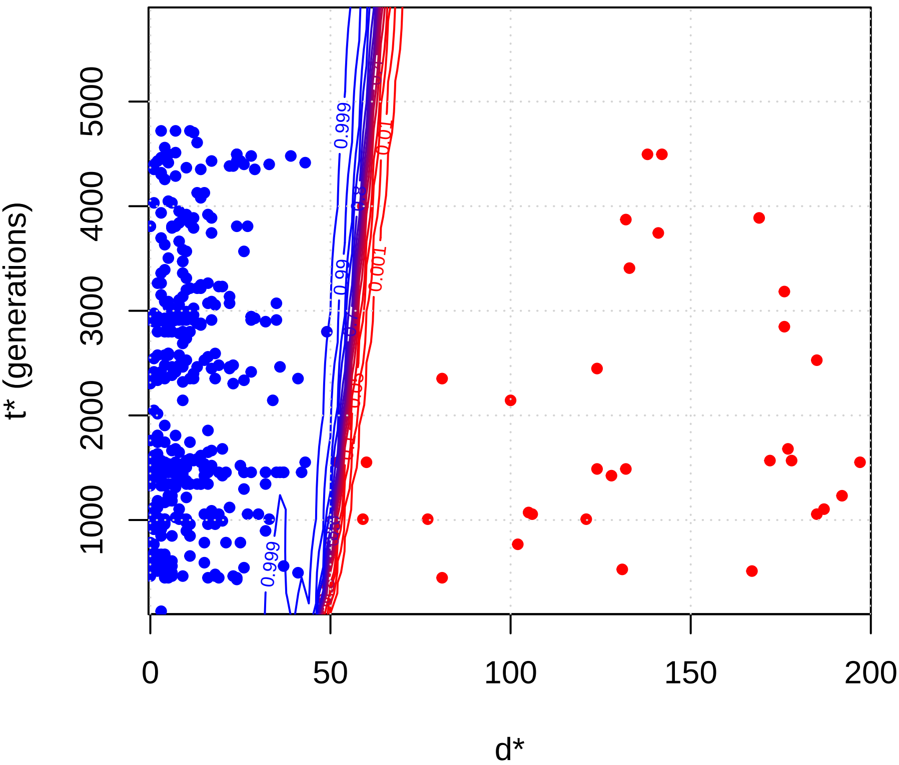

### S2 Appendix.

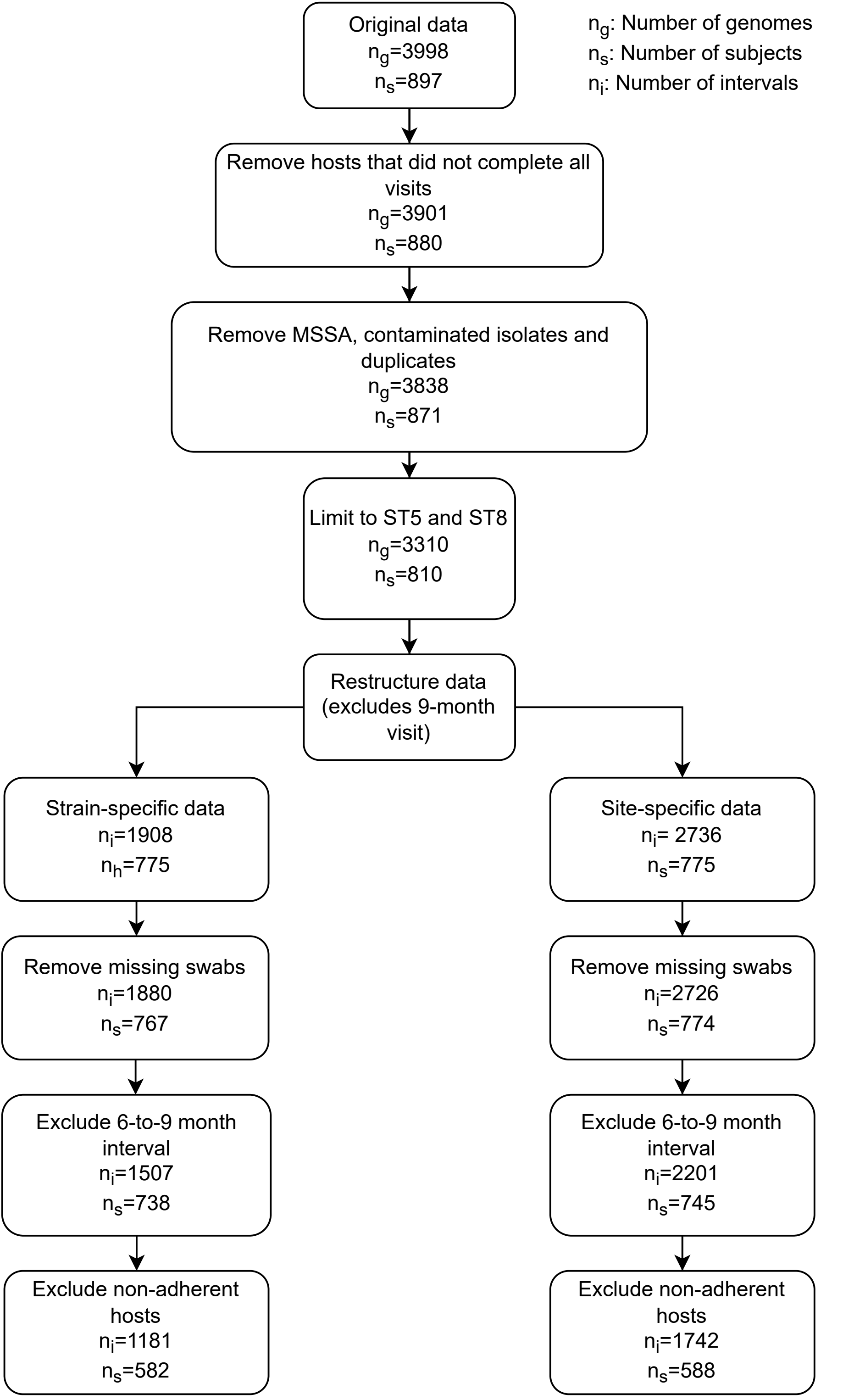

### S3 Fig.

#### Impact of resistance on MRSA clearance, Strain

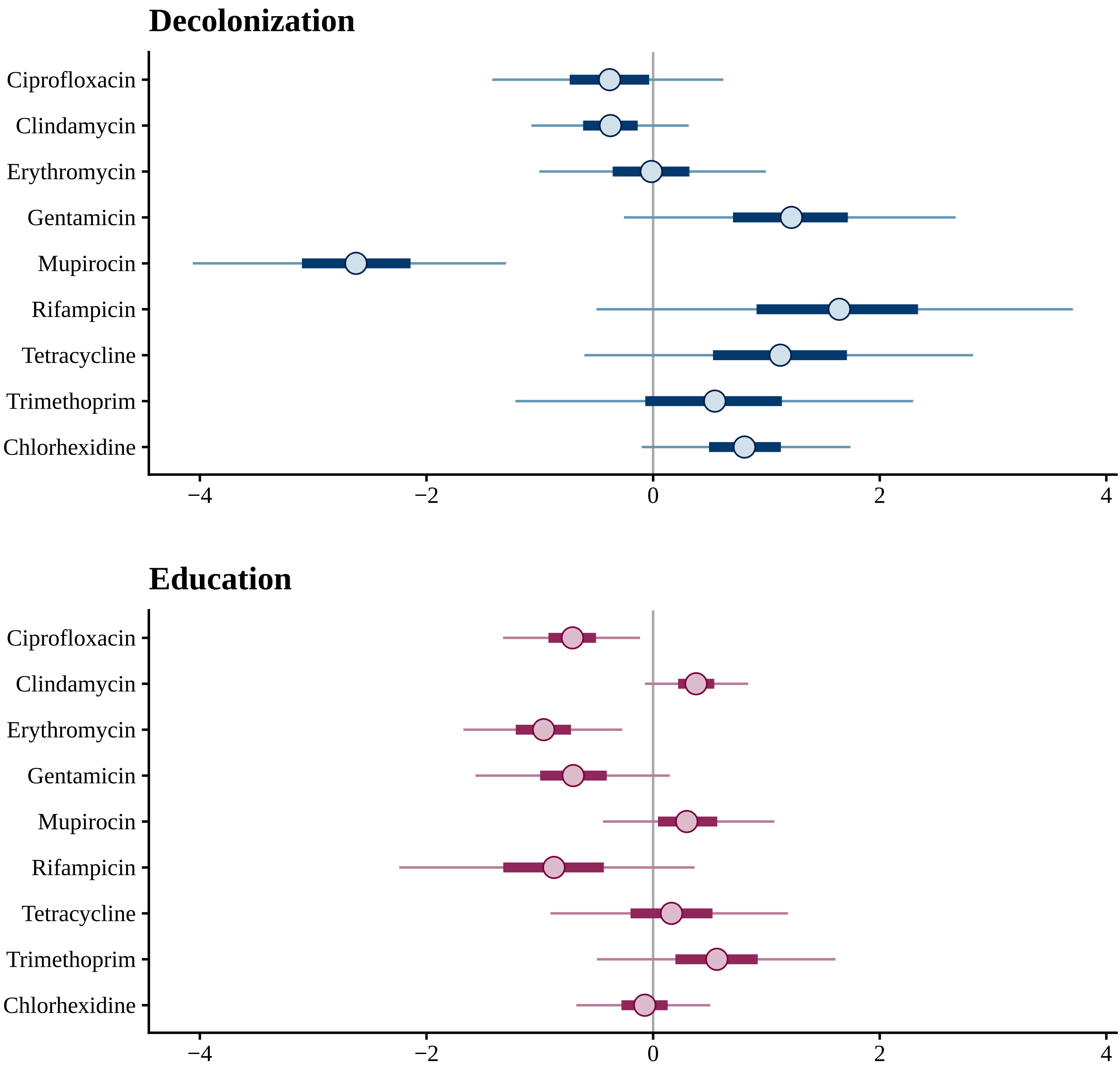

### S4 Fig.

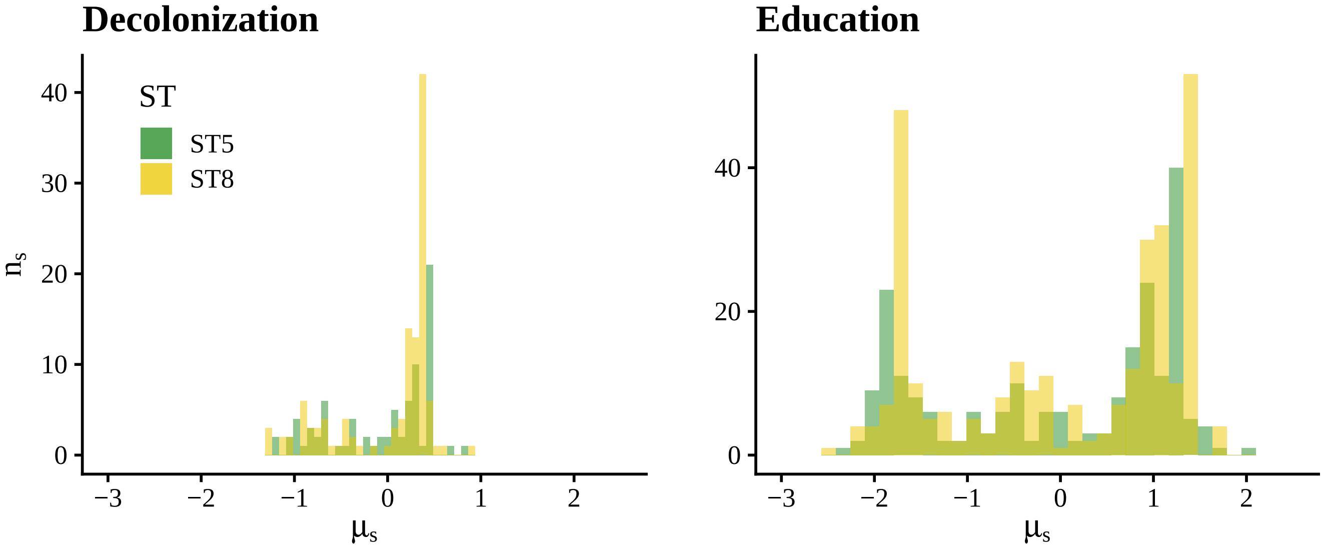

### S5 Fig.

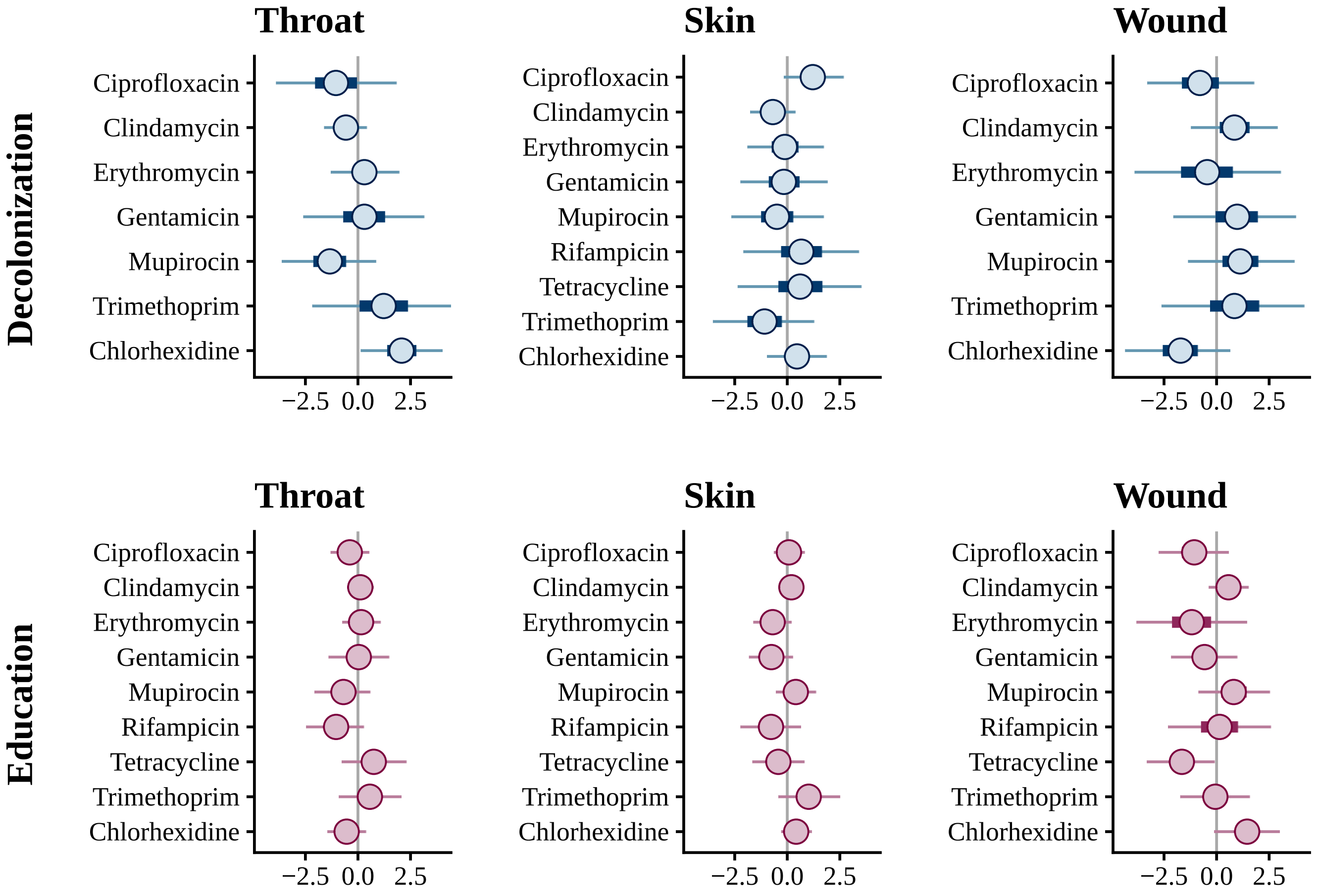

### S6 Fig.

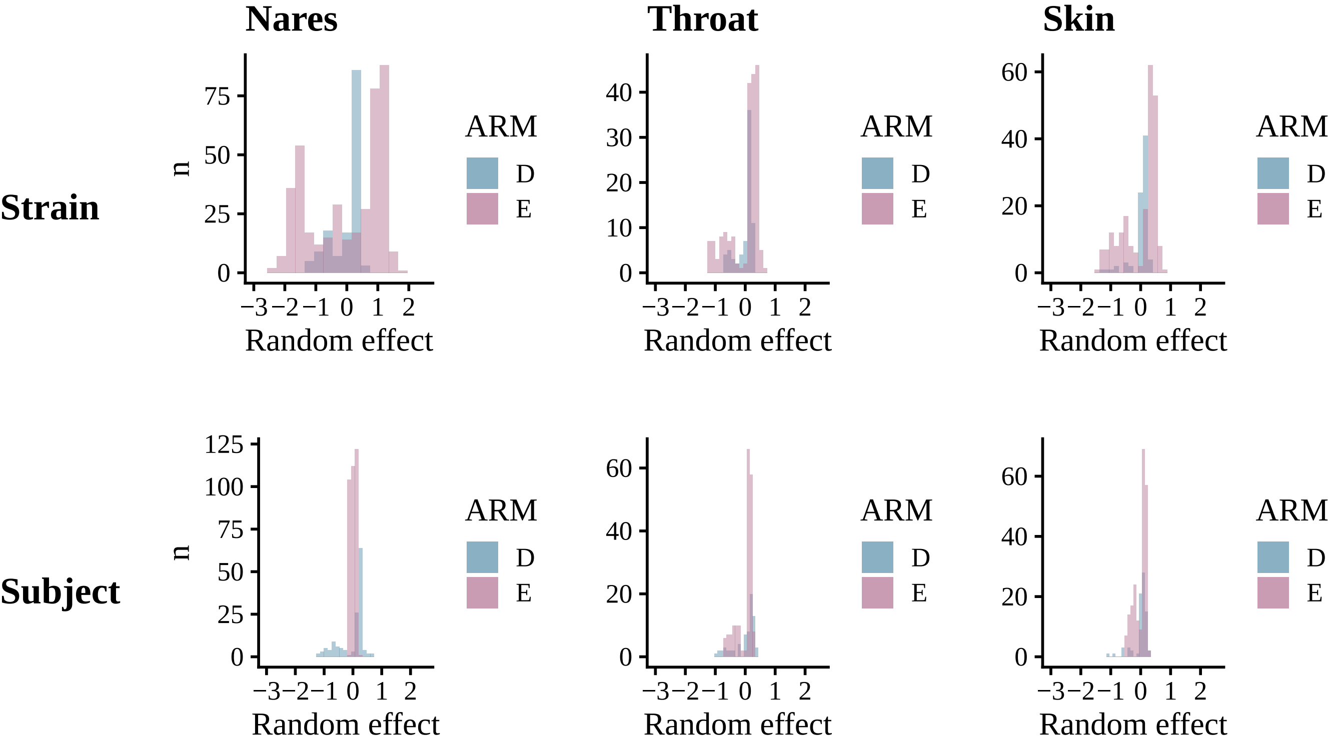
